# Universal Receptive System as a novel regulator of transcriptomic activity of *Staphylococcus aureus*

**DOI:** 10.1101/2024.09.11.612522

**Authors:** George Tetz, Kristina Kardava, Maria Vecherkovskaya, Alireza Khodadadi-Jamayran, Aristotelis Tsirigos, Victor Tetz

## Abstract

Our previous studies revealed the existence of a Universal Receptive System that regulates interactions between cells and their environment. This system is composed of DNA- and RNA-based Teazeled receptors (TezRs) found on the surface of prokaryotic and eukaryotic cells, as well as integrases and recombinases.. In the current study, we aimed to provide further insight into the regulatory role of TezR and its loss in *Staphylococcus aureus* gene transcription. To this end, transcriptomic analysis of *S. aureus* MSSA VT209 was performed following the destruction of TezRs. Bacterial RNA samples were extracted from nuclease-treated and untreated *S. aureus* MSSA VT209. After destruction of the DNA-based-, RNA-, or combined DNA- and RNA-based TezRs of *S. aureus*, 103, 150, and 93 genes were significantly differently expressed, respectively. The analysis revealed differential clustering of gene expression following the loss of different TezRs, highlighting individual cellular responses following the loss of DNA- and RNA-based TezRs. KEGG pathway gene enrichment analysis revealed that the most upregulated pathways following TezR inactivation included those related to energy metabolism, cell wall metabolism, and secretion systems. Some of the genetic pathways were related to the inhibition of biofilm formation and increased antibiotic resistance, and we confirmed this at the phenotypic level using *in vitro* studies. The results of this study add another line of evidence that the Universal Receptive System plays an important role in cell regulation, including cell responses to the environmental factors of clinically important pathogens, and that nucleic acid-based TezRs are functionally active parts of the extrabiome.

## 1 Introduction

Bacterial adaptation is a critical component of survival in changing environments and was believed to be solely regulated by protein-based receptors ((1–4)). Recently, we discovered a Universal Receptive System in both prokaryotes and eukaryotes that is composed of certain types of extracellular DNA and RNA molecules possessing receptive and regulatory properties, and also involves integrases and recombinases (5,6). This discovery sheds light on a novel aspect of cellular reception and regulation and indicates that the mechanisms involved are more complex than previously understood. These DNA- and RNA-based receptors located outside the cell wall, called Teazeled receptors (TezRs), form an extracellular network that plays an important role in cell maintenance and regulates the interactions of the cell with a diverse array of environmental factors (5,6). The Universal Receptive System has been shown not only to orchestrate the work of known protein-based receptors but also to regulate the response of cells to factors whose reception and regulation were previously unknown, including light, geomagnetic fields, and temperature. Moreover, the Universal Receptive System contributes to the formation, maintenance, and “erasure” of cellular memory (5,6). These results also echo another research study in, where we recently demonstrated previously unknown functions of DNA and RNA molecules, showing that extracellular nucleic acids act as “pliers” that alter the conformation of already synthesized proteins (7–9).

Although the role of the Universal Receptive System has been revealed in the regulation of the growth of different Gram-positive and Gram-negative bacteria, the underlying transcriptomic alterations behind these alterations remain largely unknown. The only in-depth analysis of transcriptomic alterations following the inactivation of TezRs was performed in *Bacillus pumilus*, which revealed their role in an unusual and rapid bacterial directional migration to large distances along with the control of certain virulence factors (10,11). These results suggest that TezRs are involved in complex regulatory networks that govern bacterial behavior and pathogenic potential.

Here, we analyzed the results of the alteration of the Universal Receptive System following TezR inactivation in the pathogen *Staphylococcus aureus*. *S. aureus* has been implicated in a wide range of infections characterized by its severity, immune resistance, and high rate of antibiotic resistance (12). Unlike many bacterial pathogens that possess only one or a few virulence factors, *S. aureus* produces a diverse array of virulence factors, the expression of which is predominantly regulated by two-component regulatory systems and a family of DNA-binding proteins (11,13).

Another feature of *S. aureus* infection that makes treatment particularly difficult is the high frequency of antibiotic resistance (14). *S. aureus* isolates resistant to cell wall-inhibitory antibiotics, such as β-lactam and glycopeptide antibiotics known as methicillin-resistant or vancomycin-resistant represent a significant clinical challenge.

*S. aureus* responds to cell wall antimicrobials using the two-component regulatory system, VraSR which upregulates a set of genes involved in cell-wall biosynthesis, repair, and stress (15,16). In response to beta-lactam exposure, the system triggers the production of penicillin-binding proteins, which continue to synthesize peptidoglycans despite the presence of beta-lactams (16). For vancomycin, the system induces the expression of genes that modify the d-Ala-d-Ala termini of peptidoglycan precursors, reducing vancomycin binding affinity and helping the bacterium counteract the effects of antimicrobial agents. Accordingly, mutations or downregulation of VraS and VraR decrease the development of resistance to these antibiotics (15,17).

Along with antibiotic resistance, *S. aureus* is characterized by increased tolerance to the negative effects of the outer environment and antibiotic therapy due to the formation of biofilms in which, compared to planktonic growing bacteria, they are protected from the outer environment with additional membrane-like structure and extracellular polymeric substance (18–20). Upon biofilm maturation, the bacteria within these microbial communities are up to 1,000 less sensitive to antibiotics (21,22). Biofilm formation is a sequential process regulated by the expression of different genes that govern bacterial attachment, biofilm structure development, maturation, and dispersal (23). However, the regulation of biofilm formation remains insufficiently understood despite its significant medical importance.

Here, for the first time, we conducted an in-depth functional annotation of transcriptome alterations and analysis of the alteration of Kyoto Encyclopedia of Genes and Genomes (KEGG) pathways following alteration of *S. aureus*’s Universal Receptive System by the inactivation of cell-surface-bound DNA- and RNA-based TezRs, and have shown their previously unknown role in the regulation of bacterial growth, biofilm formation, and antibiotic resistance (24).

## 2 Materials and Methods

### 2.1 Bacterial and phage strains and culture conditions

*S. aureus* MSSA VT209 was obtained from a private collection (provided by Dr. V. Tetz). Bacterial strains were passaged weekly on Columbia agar (BD Biosciences, Franklin Lakes, NJ, USA) and stored at 4 °C. All subsequent liquid subcultures were derived from colonies isolated from these plates and grown in Luria-Bertani (LB) broth (Sigma-Aldrich, St Louis, MO, USA).

### 2.2 Reagents

Human recombinant DNase I with a specific activity of 2,500 Kunitz units/mg (Sigma-Aldrich) and RNase A (Sigma-Aldrich) were used at a concentration of 10 µg/ml. Penicillin G and vancomycin (Sigma-Aldrich) were used as antibiotics.

### 2.3 Destruction of Teazled receptors from bacterial surface

To remove primary Teazled receptors (TezRs), *S. aureus* grown overnight were harvested via centrifugation at 4000 rpm for 15 min (Microfuge 20R; Beckman Coulter, La Brea, CA, USA), the pellet was washed twice in phosphate-buffered saline (PBS, pH 7.2) (Sigma-Aldrich) to an optical density at 600 nm (OD_600_) of 0.5 (Promega, GloMax, Madison, WI, USA). Bacteria were treated for 30 min at 37 °C with nucleases (DNase I or RNase A), washed three times in PBS or broth, centrifuged at 4000LJ× *g* for 15 min after each wash, and resuspended in PBS. *S. aureus*, whose TezRs were destroyed with nucleases, were marked with the superscript letter “d.” Therefore, *S. aureus* after the treatment: (i) with DNase were marked TezR–D1^d^, (ii) with RNase were marked TezR–R1^d^, and (iii) with DNase and RNase marked TezR–D1^d^/R1^d^

### 2.4 Determination of minimum inhibitory concentrations

The minimum inhibitory concentrations (MICs) of the antibiotics against nuclease-pretreated *S. aureus* were determined using the broth microdilution method according to the CLSI guidelines (25). A standard inoculum of nuclease-pretreated *S. aureus* or untreated *S. aureus* at 5 × 10^5^ colony forming units/ml (CFU)/mL was used. Serial two-fold dilutions of the antimicrobials were prepared in cation-adjusted LB broth. Bacteria were cultivated with antibiotics for 4 or 8 h, after which the MIC was defined as the lowest concentration of antibiotic that completely inhibited bacterial growth, as measured at OD_600_ (Promega, GloMax) (26). All experiments were conducted in triplicates.

## 2.5 Effect of TezR destruction on *S. aureus* biofilm formation

In each well of a 96-well flat-bottom microtiter plate (Falcon, Corning, Durham, USA), 200 μL of a standardized *S. aureus* inoculum (5 × 10^5^ CFU/mL in LB) were added. Post-nuclease treatment *S. aureus* were generated as described above. *S. aureus* without nuclease treatment was used as a control. Following 8 h incubation at 37 °C, biofilm sample well contents were aspirated and each well was washed thrice using 200 μL PBS (Sigma-Aldrich). Subsequently, 100 μL of 0.1% crystal violet (Innovating Science, Aldon corporation, Avon, USA) solution was added to the wells with dried biofilms. After 15 min, excess crystal violet was removed, wells were washed thrice with sterile water, and 150 μL of 95% ethanol was added. The absorbance was measured at 570 nm (Promega, GloMax). The tests were performed in triplicate in three separate experiments.

### 2.6 Generation of RNA sequencing data

To isolate RNA, the *S. aureus* suspension obtained 2.5 h post-nuclease treatment were washed thrice in PBS (Sigma-Aldrich) and centrifuged each time at 4000LJ× *g* for 15 min (Microfuge 20R, Beckman Coulter) followed by resuspension in PBS.

The RNeasy Mini Kit (Qiagen) was used to isolate RNA, according to the manufacturer’s instructions. The quality of the RNA was spectrophotometrically evaluated by measuring the UV absorbance at 230/260/280 nm using a NanoDrop OneC spectrophotometer (Thermo Fisher Scientific, Waltham, MA, USA).

Ribodepletion was performed using the Ribo-Zero Magnetic Gold kit (Epicenter, Madison, WI, USA) according to the manufacturer’s instructions. RNA-seq libraries were prepared using the Illumina TruSeq Stranded Total RNA Library Prep Kit. RNA libraries were pooled and sequenced using a 2 × 150 nucleotide paired-end strategy (Illumina NextSeq 500, Illumina, San Diego, CA, USA) (130 MM max).

### 2.7 RNA sequencing data processing

Sequencing reads were mapped to the reference genome of S. aureus NCTC 8325 (NCBI Reference Sequence: NC–007795) using the Bowtie2 (v2.2.4) (PMID: 22388286) and expression levels were estimated using Geneious 11.1.5. Transcripts with an adjusted p value of < 0.05 and log2 fold change value of ± 0.5 were considered for significant differential expression. The read count tables were generated using HTSeq (v0.6.0) (27), normalized based on their library size factors using DEseq2 and differential expression analysis was performed (28). To compare the level of similarity among the samples and their replicates, we used two methods: principal-component analysis and Euclidean distance-based sample clustering. All the downstream statistical analyses and generating plots were performed in R environment (v3.1.1) (https://www.r-project.org/).

We performed Kyoto Encyclopedia of Genes and Genomes (KEGG) pathway analysis for DEGs to find out their lurking functions by using R package “clusterProfiler.” (29) Gene sets at p < 0.05 were considered significantly enriched (24). Gene set enrichment analysis was performed using GSEA tool (30). Venn diagram was generated by using previously published tools (31).

### 2.8 Statistics

At least three biological replicates were performed for each experimental condition, unless stated otherwise. Each data point was denoted by the mean value ± standard deviation (SD). A two-tailed *t*-test was performed for pairwise comparisons, and statistical significance was set at p < 0.05. Statistical analyses for the biofilm assay were performed using Student’s *t*-test. GraphPad Prism version 10 (GraphPad Software, San Diego, CA, USA) or Excel 11 (Microsoft, Redmond, WA, USA) was used for statistical analyses and illustrations.

## 3 Results and Discussion

### 3.1 General Features of the transcriptome profile following TezR destruction

To analyze the consequences of the modulation of the Universal Receptive System in *S. aureus*, we performed RNA-seq analyses 2.5 h after the loss of cell surface-bound DNA-(TezR-D1), RNA-(TezR-R1), or combined DNA- and RNA-based (TezR-D1/R1) TezRs. Analysis of differentially expressed genes (DEGs) using adjusted log2 fold change > 0.5 and P-value < 0.05 revealed different patterns of gene expression following the loss of different primary TezRs (32). There were 103 DEGs with 44 and 59 genes showing up- and downregulation of *S. aureus* following the loss of TezR-D1, 150 DEGs with 137 and 13 genes up- and downregulated for *S. aureus* after the destruction of TezR-R1, and 93 DEGs with 32 and 61 genes up- and downregulated for *S. aureus* following the combined loss of TezR-D1 and TezR-R1, respectively, compared with untreated cells (**Supplementary Table 1**). The Venn diagram showed clear differentiation and clustering of DEGs between untreated *S. aureus* and those following the loss of TezR-D1, TezR-R1, or TezR-D1/R1 (Figure 1).

**Figure 1.**
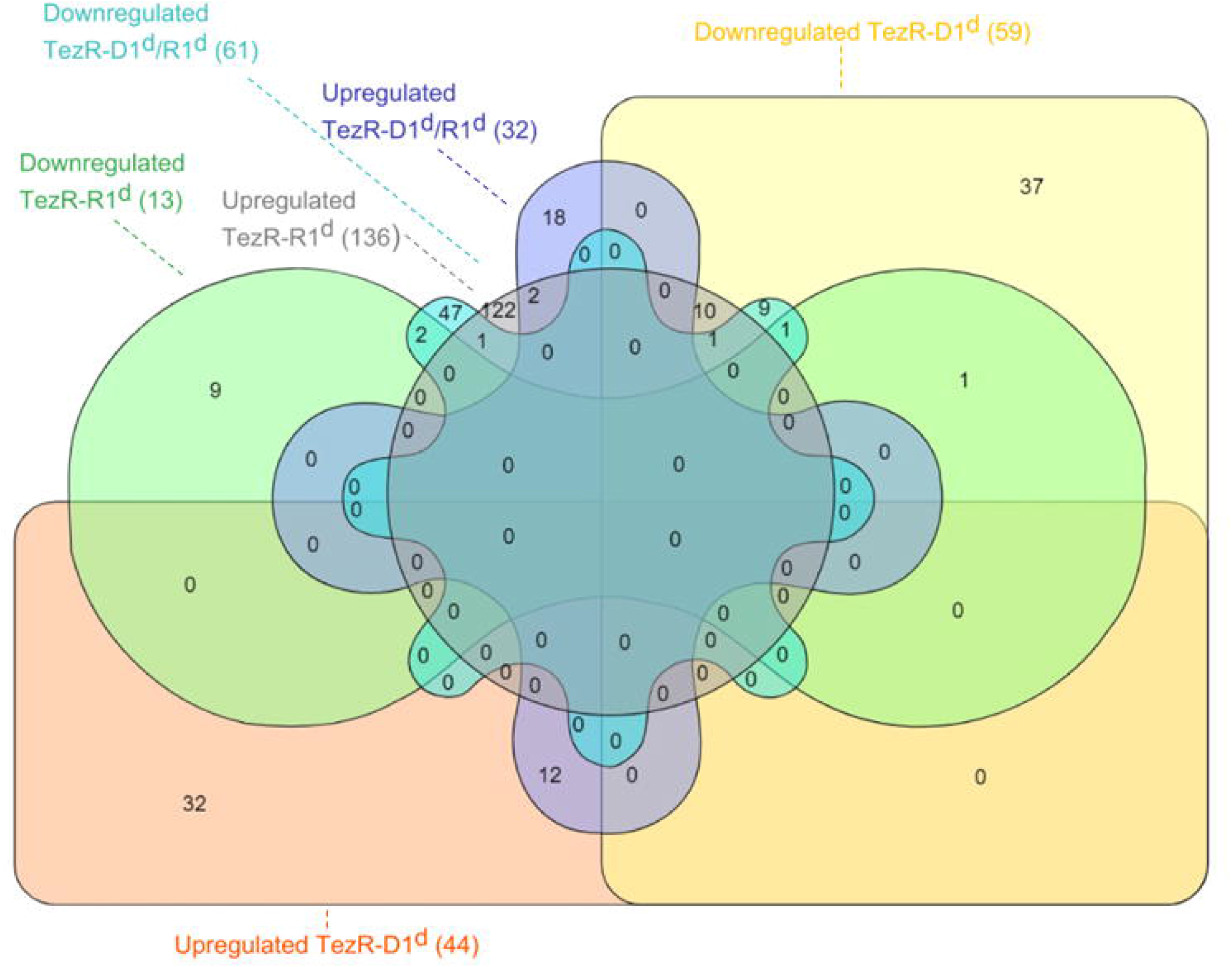
Venn diagram depicting the different regulated and overlapping genes between up- and downregulated DEGs following the destruction of DNA-based TezR (TezR–D1^d^), RNA-based TezR (TezR–R1^d^) or DNA-based and RNA-based TezR (TezR–D1^d^/R1^d^).

### 3.2 Differently expressed gene analyses following TezR-D1 destruction

The loss of DNA-based TezRs (TezR-D1) resulted in the upregulation of 44 DEGs and downregulation of 59 DEGs (Supplementary table 1). To analyze how the destruction of TezR-D1 affected *S. aureus* pathways, we mapped these DEGs to the KEGG database and analyzed them using KEGG pathway enrichment analysis. Among the top-3 KEGG pathways of the upregulated DEGs with the highest enrichment factors, two represented amino acid metabolism and included the histidine and arginine biosynthesis pathways, and the third represented the secretion system pathway (Figure 2, Supplementary Table 2).

**Figure 2.**
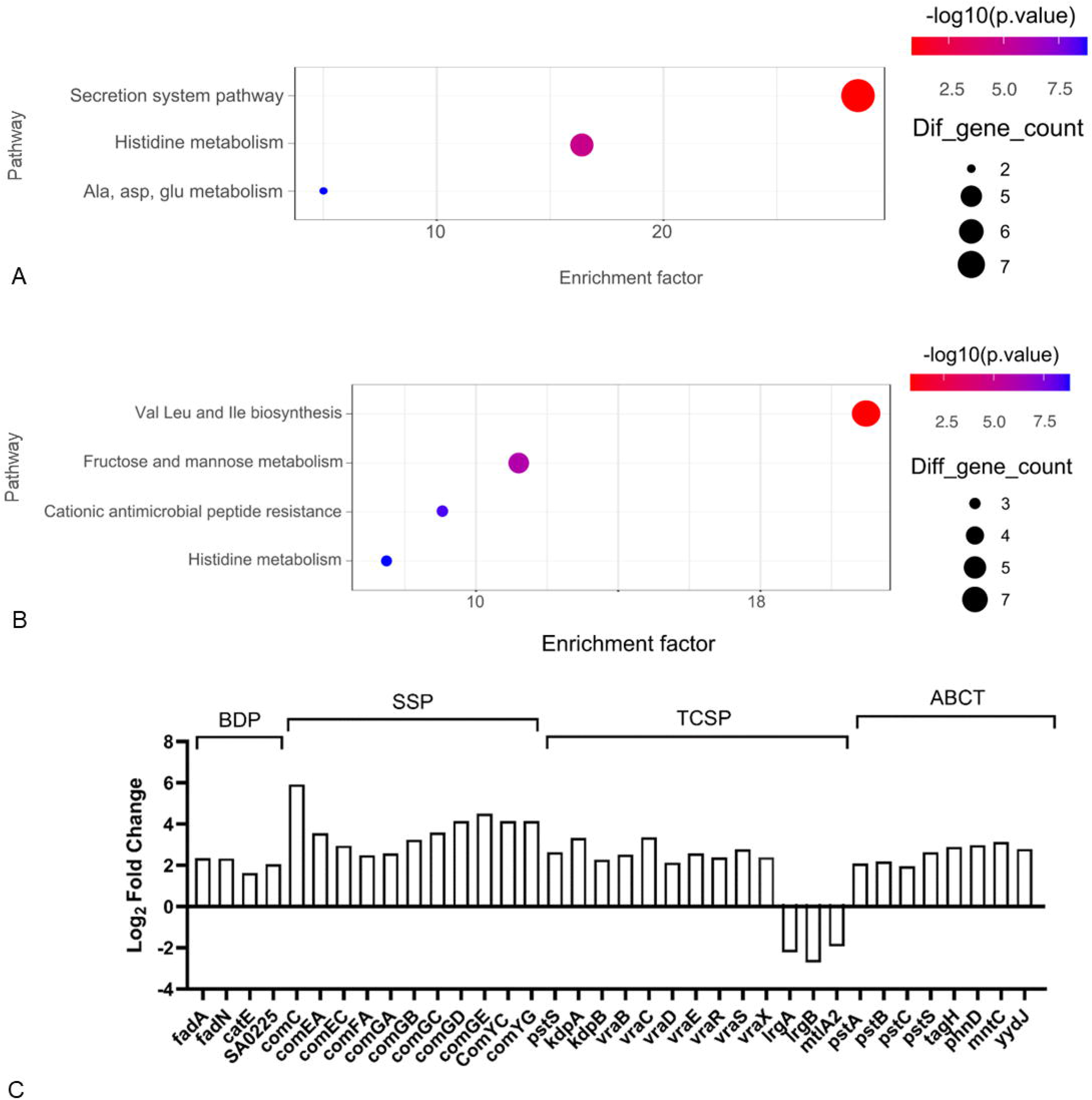
Effect on gene expression of *S. aureus* following TezR-D1 destruction. KEGG pathway enrichment of (A) up-regulated DEGs and (B) downregulated DEGs. Each circle in the graph represents a KEGG pathway, with its name in the Y-axis and the enrichment factor indicated in the X-axis. Higher enrichment factor means a more significant enrichment of the DEGs in a given pathway. The color of the circle represented the p-value. The sizes of the circles represent the number of enriched genes. The enrichment factor was defined as follows: (Number of DEGs in a term/total number of DEGs)/(total number of genes in the database in a term/total number of genes in the database). The term ‘diff gene count’ refers to the number of DEGs enriched in a KEGG pathway. (C) Analysis of differentially expressed genes (DEGs) (log2fold change > 0.5; p < 0.05) in top-3 upregulated and top-3 downregulated pathways. Levels of log2fold alteration of the expression involved in histidine metabolism pathway (HMP), aspartate and glutamate-metabolism pathway (AGP), secretion system pathway (SSP), valine, leucine and isoleucine biosynthesis pathway (VLP), Fructose and mannose metabolism pathway (FMP), and cationic antimicrobial peptide resistance pathway (CAMP).

In the histidine metabolism pathway, the enzymes involved in histidine biosynthesis, encoded by *hisA* [EC:5.3.1.16], *hisC* [EC:2.6.1.9], *hisD* [EC:1.1.1.23], *hisF* [EC:4.3.2.10], and *hisH* [EC:4.3.2.10], were significantly upregulated compared with the control (Figures 2, 3). Notably, three genes related to histidine catabolism to l-glutamate, hutU [EC: 4.2.1.49], hutI [EC: 3.5.2.7], and aldA [EC: 1.2.1.3], were downregulated, highlighting the role of this pathway in *S. aureus* adaptation following the destruction of TezR-D1.

**Figure 3.**
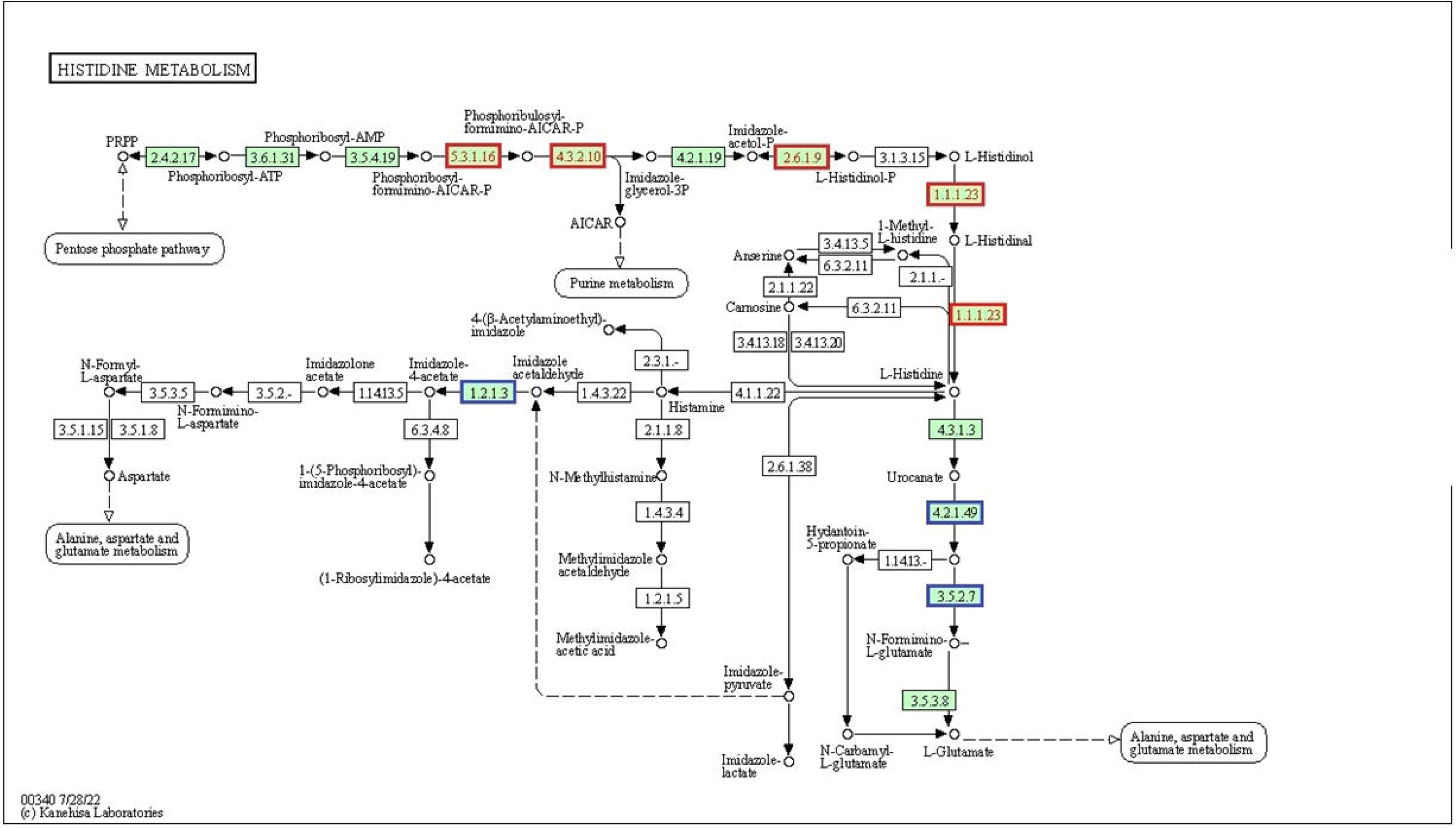
Significantly enriched KEGG pathway “Histidine metabolism” (from KEGG database). Red frames represent up-regulation and blue rectangles down-regulation of the function.

Histidine, an essential amino acid in *S. aureus*, regulates a wide range of cellular processes and cell divisions, including the formation and repair of the cell wall and cell septum formation (33–35). Therefore, the overall increase in the expression of five out of 15 key enzymes of the *S. aureus* histidine biosynthesis pathway following the loss of TezR-D1 in *S. aureus* indicates that the destruction of these cell-surface-bound DNA molecules triggers processes that require cell wall remodeling.

As shown in Figure 2A and C, along with the upregulation of histidine biosynthesis; arginine biosynthesis, involved in the TCA and urea cycles, was also upregulated upon the destruction of TezR-D1. Both *argG* and *argH*, which control the synthesis of L-arginine, an important energy source and building block for protein synthesis from glutamate or proline via citrulline metabolism, were upregulated (36).

In the “secretion system pathway” we identified seven upregulated DEGs related to type 7 secretion system (T7SS). Because the KEGG database lacks T7SS, we analyzed this pathway manually by adding genes related to T7SS, as described in Materials and Methods.

The loss of TezR-D1 resulted in the upregulation of almost the entire cluster of genes encoding the T7SS found at the ess (ESAT-6-like secretion system) locus ((37)). Therefore, along with the upregulation of three secreted proteins, EsxA, EsxB, and EsxC, which are known to play important roles in *S. aureus* pathogenicity, other integral membrane proteins, EssA, EssB, EssC, and EsaA, which are required for the synthesis and secretion of these extracellular factors, were also upregulated ((38–40)). Secreted EsxA and EsxB are pivotal for *S. aureus* virulence and persistence, modulating cytokine production and delaying the apoptosis of *S. aureus*-infected immune and epithelial cells ((38)).

Another upregulated virulence factor not revealed in the KEGG pathways was the enterotoxin Yent2, which encodes a protein that is the causative agent of toxic shock ((41,42) (Supplementary Table 2).

The highest enrichment factors of the top KEGG pathways of the downregulated DEGs are shown in Figure 2 B, С (Supplementary Table 2). Among them, the highest enrichment factors were attributed to valine, leucine, and isoleucine biosynthesis pathways, including the downregulation of genes encoding leuA [EC:2.3.3.13], leuB [EC:1.1.1.85], leuC [EC:4.2.1.33 4.2.1.35], leuD [EC:4.2.1.33 4.2.1.35], ilvA [EC:4.3.1.19], ilvB [EC:2.2.1.6], and ilvC; [EC:1.1.1.86] (Figure 4).

**Figure 4.**
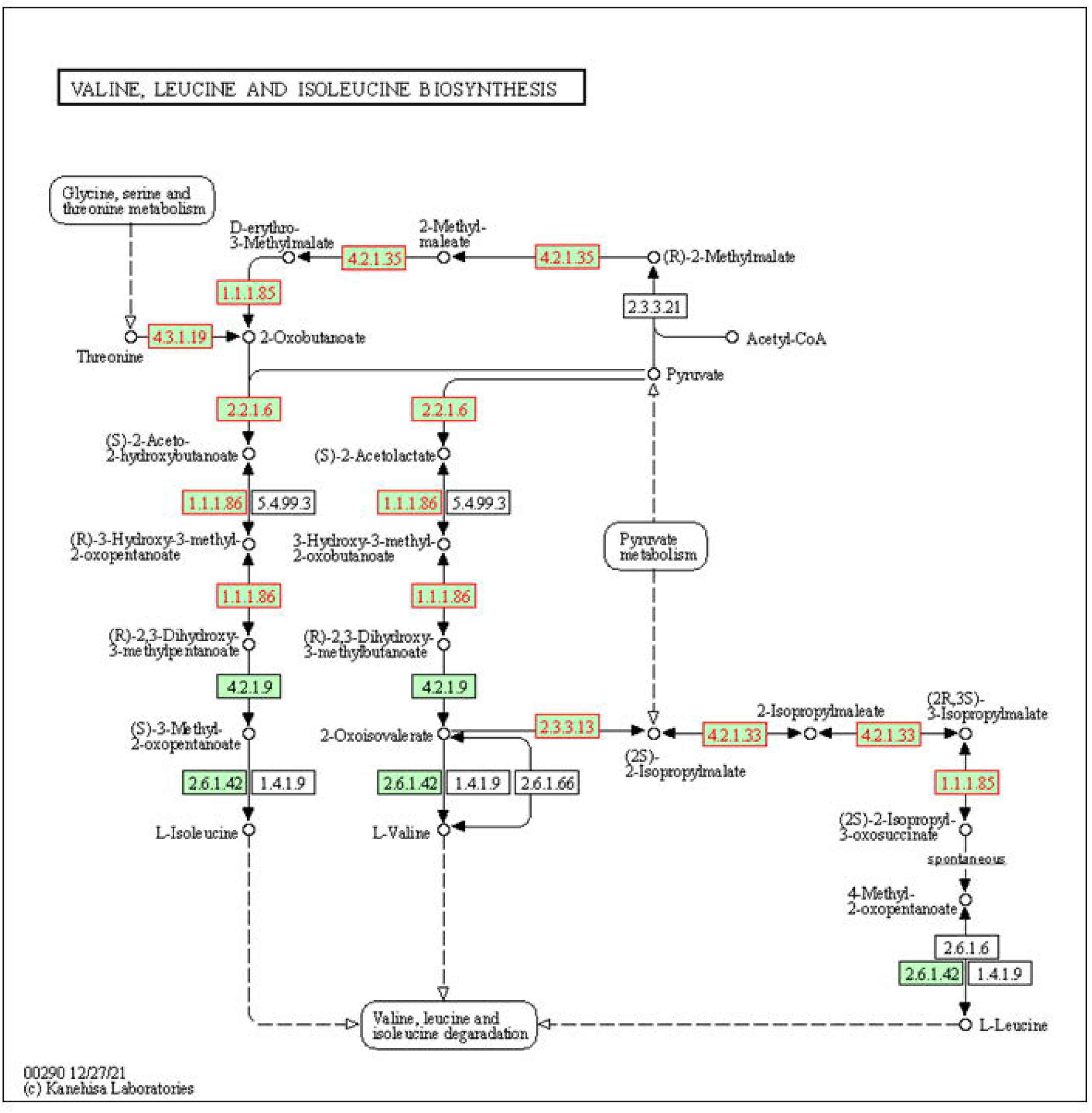
Significantly enriched KEGG pathway “Valine, leucine, and isoleucine biosynthesis pathway” (from KEGG database). Red frames represent downregulation of the function.

In *S. aureus*, branched-chain amino acids (valine, leucine, and isoleucine) represent an important group of nutrients required not only for normal metabolism and virulence but are also implicated in the synthesis of proteins and membrane branched-chain fatty acids, which play a role in membrane homeostasis and cellular adaptation (39, 40).

The fructose and mannose metabolic pathways were also enriched with downregulated enzymes encoding ManA, MtlD, ManP, MtlF, and MtlA. In Gram-positive bacteria, the ManA protein, which is a component of the mannose phosphotransferase system, participates in cell wall formation and maintains the correct carbohydrate composition of the bacterial cell wall, including teichoic acid constituents. Bacteria with depleted ManA levels are characterized by an altered cell wall architecture (44).

Following the loss of TezR-D1, three out of four genes that comprise the transcriptional regulator, and mannitol-1-phosphate dehydrogenase including *mtlA* (enzyme IICB^mtl^), *mtlF* (enzyme IIA^mtl^), and *mtlD* were downregulated. The impaired ability of *S. aureus* to assimilate mannitol plays an important role in cell adaptation through the regulation of glycolytic pathways and the maintenance of cellular redox and osmotic potential (45,46).

Finally, the pathway related to resistance to cationic antimicrobial peptide (CAMP) resistance was downregulated. We observed downregulation of the whole *dlt* operon, which comprises four genes (*dltA, dltB, dltC*, and *dltD*), although the expression of *dltC* was above the threshold of log2fold change > 0.5 (Figure 2B, Supplementary Table 1). The *dlt* operon catalyzes the incorporation of d-alanine residues into lipoteichoic acids, and the inhibition of this process in gram-positive cocci results in increased electronegativity of the bacterial surface (47). This leads to more efficient binding of cationic compounds and makes cells more susceptible to CAMPs (48,49). This finding is particularly interesting because following the destruction of negatively charged DNA-based TezRs, *S. aureus* should have higher electropositivity; therefore, the increased electronegativity by the regulation of the *dlt* operon might help maintain the cell surface charge.

### 3.3 Differently expressed gene analyses following TezR-R1 destruction

The loss of RNA-based TezRs (TezR-R1) resulted in the upregulation of 137 DEGs and downregulation of 13 DEGs (Supplementary Table 1). The upregulated genes outnumbered the downregulated genes, suggesting that gene metabolism in *S. aureus* increased following TezR-R1 loss.

The top KEGG pathways of the upregulated DEGs with the highest enrichment factors following TezR-R1 destruction were the benzoate degradation pathway, secretion system (Type II secretion system) pathway, two-component system pathway, and ABC transporters (Figure 5, Supplementary Table 3).

**Figure 5.**
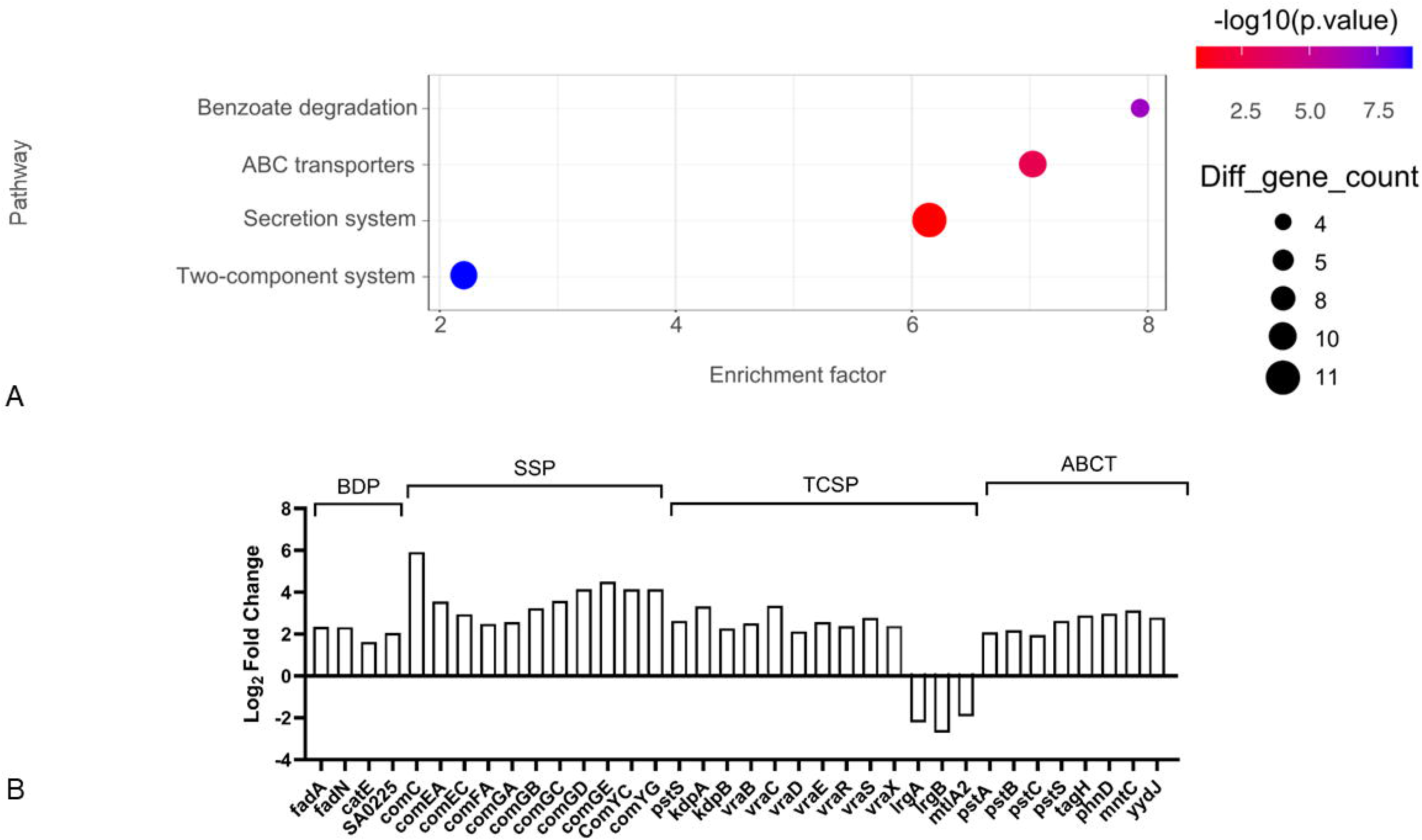
Effect on gene expression of *S. aureus* following TezR-R1 destruction. (A) KEGG pathway enrichment of up-regulated DEGs. Each circle in the graph represents a KEGG pathway, with its name in the Y-axis and the enrichment factor indicated in the X-axis. Higher enrichment factor means a more significant enrichment of the DEGs in a given pathway. The color of the circle represented the p-value. The sizes of the circles represent the number of enriched genes. The enrichment factor was defined as follows: (Number of DEGs in a term/total number of DEGs)/(total number of genes in the database in a term/total number of genes in the database). The term ‘diff gene count’ refers to the number of DEGs enriched in a KEGG pathway. (B) Analysis of differentially expressed genes (DEGs) (log2fold change > 0.5; p < 0.05) in top upregulated and downregulated pathways Levels of log2fold alteration of the expression involved in benzoate degradation pathway (BDP), secretion system pathway (SSP), two component system pathway (TCSP), and ABC transporters (ABCT).

Genes in the benzoate degradation pathway, which resulted in a higher yield of acetyl-coenzyme A, including *fadA*, *fadN*, *catE*, SA0225, and *vraB*, were upregulated (50) (Figure 5).

Notably, the upregulation of SA0225 and *fadN* is involved in the anaerobic pathway of benzoate degradation in bacteria, despite the fact that in our study, *S. aureus* was cultivated under aerobic conditions. It is consistent with recently published data showing that the loss of RNA-based TezR in aerobic-growing *Bacillus pumilus* results in the upregulation of core anaerobic energy metabolism enzymes (50–52). This echoes with the results from another study demonstrated the obligate aerobe *Pseudomonas putida* became capable of growing anaerobically after RNA-based TezR destruction (6).

Following the loss of RNA-based TezRs TezR-R1, we also observed the upregulation of genes related to the type II secretion system (T2S), which in Gram-positive bacteria is involved in transformation and acts as a machinery for DNA uptake (53). The key upregulated genes were competence proteins required for dsDNA and ssDNA to pass through the cell wall and cytoplasmic membrane, including ComEA, ComEC, comFA, ComYC, ComYG, and ComC (54,55). In addition, the genes of the *comG* operon, which are required for DNA binding to the cell surface, priming pilus assembly, and transformation, were upregulated (56). These overexpressed genes included *comGA*, which encodes the assembly ATPase required for initial extracellular DNA binding, and *comGB*, *comGC*, *comGE*, and *comGD*, which encode the pseudopilus and major and minor pilin proteins (57–59). These results indicated that the destruction of TezR-R1 resulted in the upregulation of almost the entire cluster of *S. aureus* genes related to bacterial transformation and extracellular DNA uptake.

Within the “Two component system” pathway, we observed the upregulation of *pstS*, *kdpA, kdpB*, *vraB*, *vraC*, *vraD*, *vraE*, *vraR*, *vraS*, and *vraX*. The upregulation of the VraSR regulatory system (vancomycin resistance-associated sensor-regulator system) is particularly interesting because it is known to lead to modifications in cell wall structure and is associated with resistance to cell wall-targeting antibiotics such as β-lactam and glycopeptide antibiotics (15,60). Overexpression of the *vraSR* is part of the cell wall stress resistance due to cell wall damage or inhibition of cell wall synthesis, indicating that the loss of RNA-bound TezR is regulated by the cell as a stress event for the cell wall (13,61).

Significant differences were found in the expression of genes involved in “ABC transporter” pathway. In particular, the expression of *pstSCAB*, an ABC family importer comprising the transmembrane channel, and ATPase, which energizes the translocation of inorganic phosphate during its limitation, was upregulated (62) (63). Other upregulated proteins related to ABC transporters and involved in the transport of manganese ions and phosphonates as well as the export of teichoic acids play a role in oxidative stress protection, including MntC, TagH, PhnD, and YydJ (64,65).

Additionally, we observed that the overexpressed DEGs related to *S. aureus* virulence were not covered by the KEGG pathways. We found upregulation of genes relevant for host-pathogen interactions, including SA0221, which blocks C3 convertases; *fmtA*, which promotes antimicrobial peptide resistance, neutrophil survival, and epithelial cell invasion; and exotoxin *set15*, which inhibits the host’s innate immune response (Supplementary Table 1) (66,67).

There were only 11 downregulated DEGs in *S. aureus* after the destruction of TezR-R1, and this number was insufficient to analyze the KEGG pathways (Supplementary table 1). However, we were particularly interested by the downregulation of *lrgA* and *lrgB* of the *lrgAB* operon at “Two component system” pathway. Both LrgA and LrgB are associated with the control of murein hydrolase activity, which is involved in cell wall growth and the separation of daughter cells, again showing an association between TezRs and the control of cell wall growth (68). Notably, under regular stress conditions, the *lrgAB* operon is upregulated, inhibiting murein hydrolase activity, and thereby preventing cell lysis in unfavorable environments (69). Another down-regulated protein was MtlA2, which is responsible for the efficient phosphorylation of mannitol and its utilization as a carbon source (70).

### 3.4 Differently expressed gene analyses following combined TezR-D1/R1 destruction

Following the combined loss of cell-surface-bound DNA and RNA TezR (TezR-D1/R1 loss), *S. aureus* was characterized by the upregulation of 32 and downregulation of 61 DEGs (Supplementary Table 1).

In previous studies, owing to the parodoxal responses to the outer environment by the cells following the loss of both DNA- and RNA-based TezRs, they were named “drunk cells.” The significantly upregulated KEGG altered pathways highlighted the involvement of upregulated DEGs in “Bacterial invasion of epithelial cells,” “Staphylococcus aureus infection,” and “Secretion system” pathways (Supplementary Table 4).

In the “Bacterial invasion of epithelial cells” pathway, both *fnbA* and *fnbB* were significantly up-regulated as comparison with the control. In previous studies, FnbA and FnbB have been demonstrated to be important adhesins involved in biofilm formation, cell adhesion, and host cell internalization (71–73) (Figure 6).

**Figure 6.**
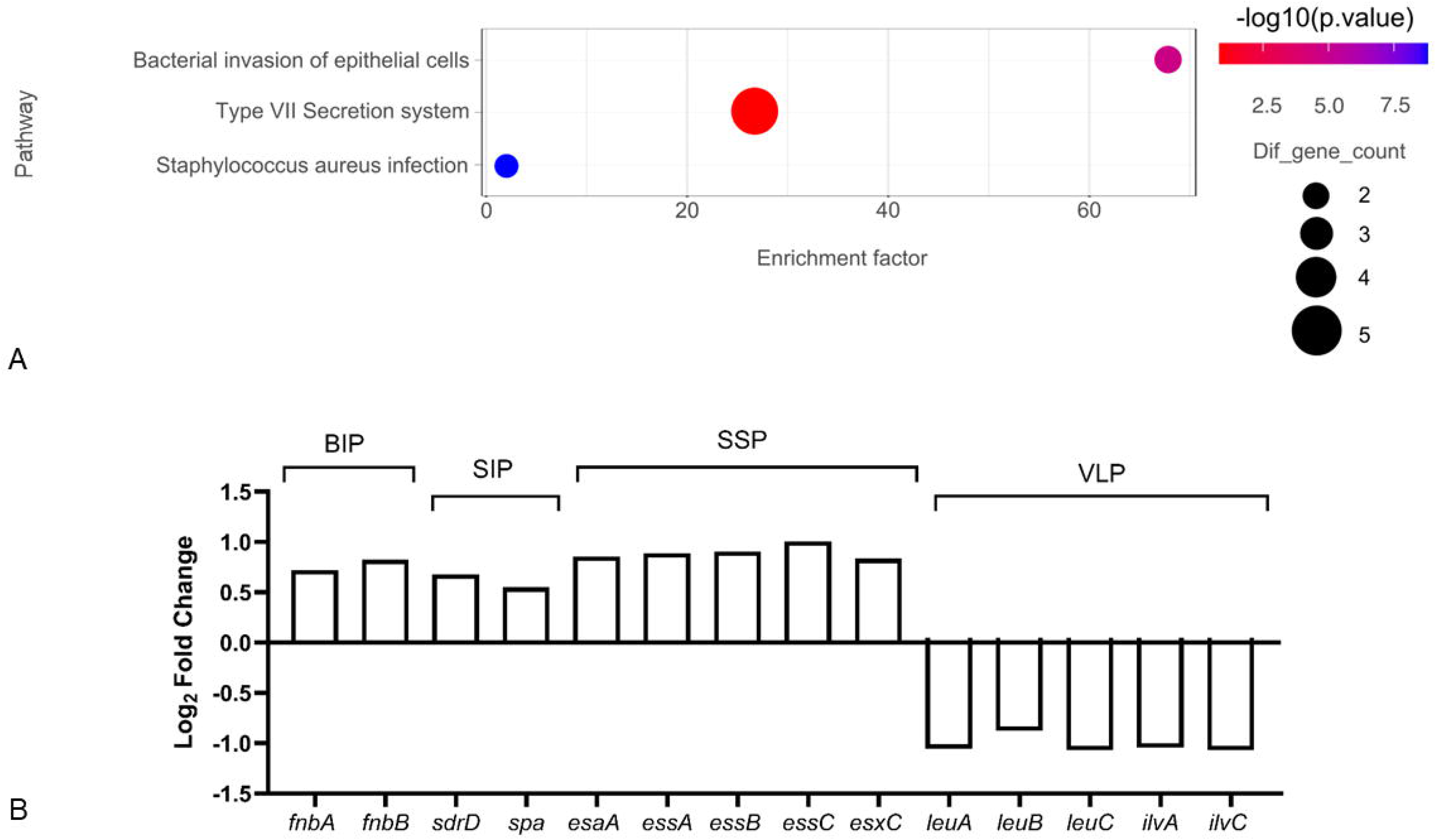
Effect on gene expression of *S. aureus* following TezR-D1/R1 destruction. (A) KEGG pathway enrichment of up-regulated DEGs. Each circle in the graph represents a KEGG pathway, with its name in the Y-axis and the enrichment factor indicated in the X-axis. Higher enrichment factor means a more significant enrichment of the DEGs in a given pathway. The color of the circle represented the p-value. The sizes of the circles represent the number of enriched genes. The enrichment factor was defined as follows: (Number of DEGs in a term/total number of DEGs)/(total number of genes in the database in a term/total number of genes in the database). The term ‘diff gene count’ refers to the number of DEGs enriched in a KEGG pathway. (B) Analysis of differentially expressed genes (DEGs) (log2fold change > 0.5; p < 0.05) in top-3 upregulated and top-3 downregulated pathways: Levels of log2fold alteration of the expression involved in benzoate degradation pathway (BDP), bacterial invasion of epithelial cells pathway (BIP), *Staphylococcus aureus* infection pathway (SIP), secretion system pathway (SSP), valine, leucine and isoleucine biosynthesis pathway (VLP).

In another upregulated infection-related pathway, the “*S. aureus* infection pathway” two genes, sdrE and spa were upregulated. The surface protein SdrD has been shown to interact directly with the complement control protein factor H, facilitating staphylococcal infection, and in some cases, is involved in platelet aggregation. Another upregulated cell surface-associated protein, Spa, promotes bacterial growth and virulence, and is also implicated in quorum sensing (74–77).

In *S. aureus* following the loss of DNA- and RNA-based TezRs, like following the isolated loss of only DNA-based TezRs, we observed upregulation of “Secretion system” pathway including T7SS genes which we manually added to the analysis due to their lack in KEGG database. The destruction of TezR-D1/R1 upregulates integral membrane proteins (EsaA, EssA, EssB, and EssC) and the secreted substrate EsxC, which are important for the persistence of *S. aureus* infection (78–80).

Among other genes that attracted our attention but were not covered by existing KEGG pathways, the methyltransferase *rlmN*, which modifies A2503 in 23S rRNA, showed the highest upregulation of over 4.5 fold (Supplementary Table 1). The role of RlmN in *S. aureus* is not fully understood, with previous studies showing its involvement in the interaction of the ribosome with the nascent peptide, thus affecting translational speed and partially controlling antibiotic susceptibility (81,82).

The analysis revealed that the only significantly enriched pathways among 61 downregulated DEGs were “Valine, leucine and isoleucine biosynthesis” (Figure 6).

The top depressed genes following the combined destruction of TezRs-D1 and TezR-R1 involved in “Valine, leucine and isoleucine biosynthesis” pathway were within ilv-leu region and included *leuA*, *leuB*, *leuC*, *ilvA*, *ilvC* controlling the synthesis of leucine and isoleucine (Figure 6). Branched-chain amino acids are critical for the synthesis of anti-branched-chain fatty acids, which play a role in membrane fluidity (83). These data open the discussion on the role of cell surface-bound DNA- and RNA-bound TezRs in membrane fluidity; however, more detailed related studies are planned to be highlighted in separate articles.

Among the other genes not covered by the KEGG pathways, which attracted our attention were downregulated genes associated with stress response-related genes, including *hupB*, *dps*, and *ftsL,* that facilitate responses to various environmental changes.

In addition, among the genes implicated in transcription, *yydK*, *cggR*, *mraZ*, *hrcA*, and *ctsR* were downregulated. Downregulation of CtsR- and HrcA-controlled heat shock regulation is particularly interesting because this downregulation is known to enhance bacterial survival under unfavorable temperatures (84).

The depletion of *cggR*, which encodes a glycolytic repressor, has also attracted our attention. CggR activates glycolysis, which is typical of *S. aureus* under anaerobic conditions (85,86). These data are in agreement with our previous findings that the Universal Receptive System plays an important role in regulating anaerobic energy metabolism pathways (6,10)(6,10). Notably, this pathway for the activation of anaerobic energy metabolism was different from that observed in *S. aureus* following the loss of RNA-based TezRs alone, highlighting the individuality of cell responses following the loss of different types of TezRs.

### 3.5 Effect of TezR loss on biofilm formation

In this study, we observed that following the loss of DNA-based TezR, some pathways related to the inhibition of biofilm formation were altered. For example, the upregulation of *argG* and *argH*, which are responsible for arginine formation, is known to inhibit biofilm formation in *S. aureus* via several pathways, including the prevention of bacterial co-aggregation. Additionally, the downregulation of numerous genes within the fructose and mannose metabolic pathways negatively impacts biofilm formation by inhibiting the transport and phosphorylation of sugars (87–90). Simultaneously, the loss of RNA-based TezRs, or combined DNA- and RNA-based TetzRs, did not alter the transcriptomic activity of known genes involved in biofilm formation.

To confirm these transcriptomic alterations following TezR-D1 loss at the phenotypic level, we analyzed the effects of TezR depletion on biofilm formation by *S. aureus* (Figure 7).We analyzed the dynamics of biofilm formation in its early phases, since we have previously shown that TezRs following loss restore their function in approximately 8 h (6).

**Figure 7.**
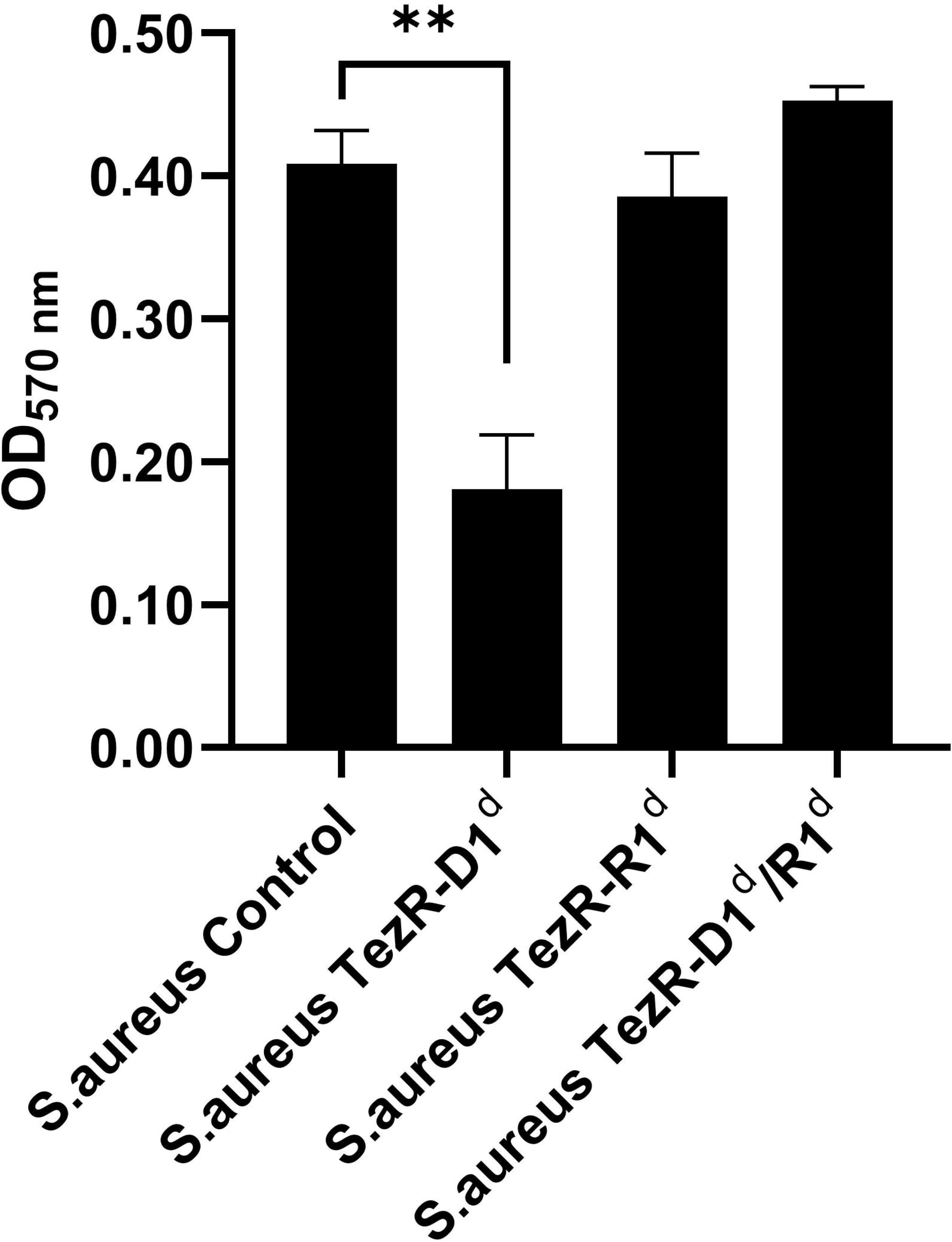
Biofilm formation of *S. aureus* following the loss of different TezRs at different time points. Statistically significant differences **p < 0.01). *S. aureus* following the loss of: DNA-based TezRs (TezR-D1^d^), RNA-based TezRs (TezR-R1^d^), DNA- and RNA-based TezRs (TezR-D1^d^/R1^d^). Data represent the meanLJ±LJSD from three independent experiments.

After 8 h, *S. aureus* biofilm exhibited OD_570_ nm values of 0.408 ± 0.019. The loss of DNA-based TezRs resulted in the inhibition of biofilm formation; thus, the biofilms formed by *S. aureus* TezR-D1^d^ exhibited OD_570_ nm values of 0.181 ± 0.031 after 8-h incubation. Loss of RNA-based and combined DNase- and RNase-based TezRs did not affect biofilm formation.

These findings are particularly interesting because numerous previous studies have shown that DNase treatment inhibits *S. aureus* biofilm formation. The data from these articles suggest that the action of this nuclease against microbial biofilms occurs through the inhibition of bacterial adhesion by destroying sticky extracellular DNA or by the inhibition of the extracellular DNA’s role as a gene messenger (91–95). However, in all of these studies, cell-surface bound DNA-based TezR was also methodologically affected by the addition of DNase, but the role of the loss of these elements in the regulation of biofilm formation seen in the transcriptomic analysis was not taken into consideration. Moreover, the lack of an effect of the combined DNase and RNase treatment observed in this study highlights the specificity of the bacterial response to isolated cell surface-bound DNA degradation, confirming the specificity of transcriptomic alterations following the loss of different types of primary TezRs.

### 3.6 Effect of TezRs loss on antibiotic resistance

Finally, we compared the correlation between transcriptomic alterations in genes related to antibiotic resistance, which were upregulated following modulation of the Universal Receptive System, and phenotypic resistance to antibiotics. According to the transcriptomic analysis, the only known DEGs directly related to antibiotic resistance were genes involved in the two-component regulatory system following the destruction of RNA-based TezRs. Specifically, we observed upregulation of the VraSR regulon, which is a key element in the generation of a highly resistant *S. aureus* phenotype associated with resistance to cell wall-targeting antibiotics (17,61,96).

Therefore, we evaluated the MIC of penicillin G as a representative of beta-lactams and vancomycin at 2 and 4 h post-TezR destruction. We intentionally selected these time frames because, as was previously shown, *S. aureus* did not restore TezRs within this period (6).

*S. aureus* following the loss of DNA-based TezR (*S. aureus* TezR–D1^d^), RNA-based TezR (*S. aureus* TezR–R1^d^) or TezR–D1 and TezR–R1 (*S. aureus* TezR–D1^d^/R1^d^). Data represent the meanLJ±LJSD from three independent experiments.

Data received indicate that following the inactivation of RNA-based TezRs, the TezR-R1^d^ exhibited a resistant phenotype to penicillin G compared to control with MIC 8 µg/mL (Figure 8A). Destruction of DNA-or the combined loss of DNA- and RNA-based TezRs did not affect the sensitivity of *S. aureus* to penicillin G.

**Figure 8.**
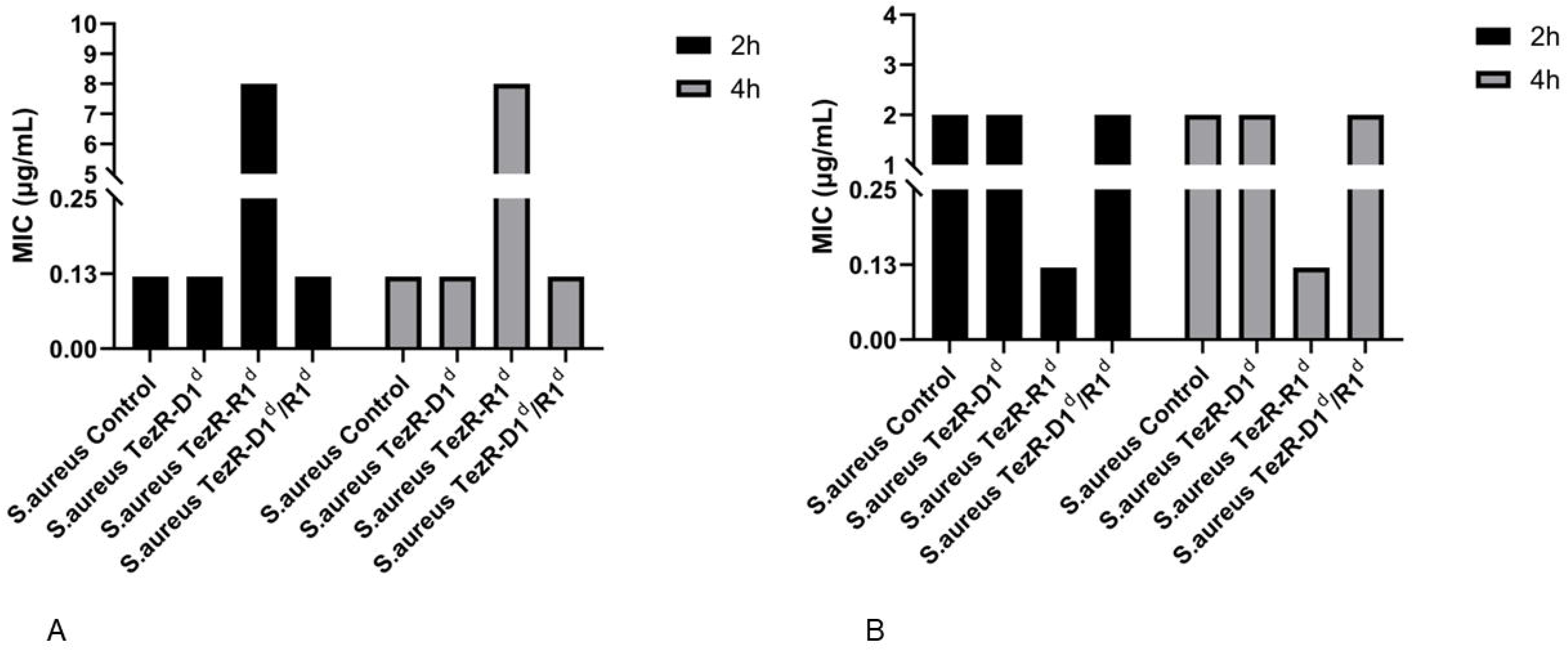
Effect of TezR loss on minimal inhibitory concentration to (A) penicillin G (B) vancomycin.

Contrary to expectations, the loss of RNA-based TezRs resulted in the higher sensitivity of *S. aureus* to vancomycin, which became 0.12 µg/mL which was lower than that in untreated control for which MIC was 0.5 µg/mL (Figure 8B). Loss of DNA- or combined DNA- and RNA-based TezR did not affect the sensitivity of *S. aureus* to vancomycin. The reason for the increased sensitivity of *S. aureus* TezR-R1^d^ to vancomycin upon upregulation of the VraSR regulon is unclear. However, it can potentially be explained by the simultaneous downregulation of *lrgB* and *lrgB*, which confers increased murein hydrolase activity owing to increased autolysis and the inability to protect against the specific mechanism of action of vancomycin on peptidoglycan precursors. These complex interplays highlight the complicated processes in bacteria controlled by the Universal Receptive System and will be studied in a separate extended study to determine their role in the sensitivity of bacteria to antibiotics (97,98).

## 4 Conclusion

This study demonstrates that the Universal Receptive system and its components, TezRs, regulate a diverse array of *S. aureus* cell activities. The proteomic profiles obtained following the loss of DNA- and RNA-based TezRs revealed changes in different KEGG pathways and altered the expression of many proteins, including those involved in the infection process and sensitivity to antimicrobial agents. Importantly, KEGG enrichment analyses of the transcriptome and proteome showed a relatively low overlap in enriched pathways following the individual or combined destruction of DNA-and RNA-based TezRs, suggesting specificity and complexity in the regulatory mechanisms controlling gene and protein expression by the Universal Receptive System. The data received add another line of evidence that the Universal Receptive System plays an important role in cell regulation, including cell responses to the environmental factors of clinically important pathogens, and that nucleic acid-based TezRs are functionally active parts of the extrabiome (99).

Future research should focus on studying the role of this system in a more diverse array of microbial pathogens, their sensitivity to antibiotics, and their implications for the regulation of real-life infections.

**Supplementary Dataset 1:** List of differentially expressed genes of *S.aureus* after nuclease treatment..

**Supplementary Dataset 2:** List of differentially expressed genes in *S. aureus* after DNase treatment used for KEGG analysis.

**Supplementary Dataset 3:** List of differentially expressed genes in *S. aureus* after RNase treatment used for KEGG analysis.

**Supplementary Dataset 4:** List of differentially expressed genes in *S. aureus* after DNase and RNase treatment used for KEGG analysis..

## 6 Data Availability Statement

The datasets for this study are available on request.

## 7 Author Contributions

V.T., G.T., K.K. designed the experiments and study. G.T., V.T., M.V. supervised data analysis and wrote the manuscript. A.K.J. conducted data analyses, A.T. gave technical support and conceptual advice.

## Acknowledgements

We would like to thank the Genome Technology Center (GTC) for expert library preparation and sequencing, and the Applied Bioinformatics Laboratories (ABL) for providing bioinformatics support and helping with the analysis and interpretation of the data. GTC and ABL are shared resources partially supported by the Cancer Center Support Grant P30CA016087 at the Laura and Isaac Perlmutter Cancer Center. This work has used computing resources at the NYU School of Medicine High Performance Computing (HPC) Facility.

