## Supplementary Table 2 for "Universal Receptive System as a novel regulator of transcriptomic activity of *Staphylococcus aureus*"

List of differentially expressed genes in *S. aureus* after DNase treatment used for KEGG analysis.

| **Locus** | **Log 2 Fold change** | **Common name** | **Description** |
| --- | --- | --- | --- |
| [SA2466](https://www.genome.jp/entry/sau:SA2466) | 1.034729 | HisA | phosphoribosylformimino-5-aminoimidazole carboxamide ribotide isomerase |
| [SA2469](https://www.genome.jp/entry/sau:SA2469) | 0.65013 | HisC | histidinol-phosphate aminotransferase |
| [SA2470](https://www.genome.jp/entry/sau:SA2470) | 0.673855 | HisD | histidinol dehydrogenase |
| [SA2465](https://www.genome.jp/entry/sau:SA2465) | 0.851572 | HisF | imidazole glycerol-phosphate synthase subunit HisF |
| [SA2467](https://www.genome.jp/entry/sau:SA2467) | 0.729793 | HisH | imidazole glycerol-phosphate synthase subunit HisH |
| [SA2121](https://www.genome.jp/entry/sau:SA2121) | -0.62255 | HutI | imidazolonepropionase |
| [SA2122](https://www.genome.jp/entry/sau:SA2122) | -0.61493 | HutU | urocanate hydratase |
| [SA0162](https://www.genome.jp/entry/sau:SA0162) | -0.71756 | AdlA | aldehyde dehydrogenase (NAD+) |
| [SA0822](https://www.genome.jp/entry/sau:SA0822) | 0.773082 | ArgG | argininosuccinate synthase |
| [SA0821](https://www.genome.jp/entry/sau:SA0821) | 0.983391 | ArgH | argininosuccinate lyase |
| SA0271 | 0.665833 | EsxA | Type VII secretion system extracellular protein A |
| SA0278 | 0.736594 | EsxB | Type VII secretion system extracellular protein B |
| SA0277 | 0.680881 | EsxC | ESAT-6 secretion system extracellular protein C |
| SA0273 | 0.680721 | EssA | ESAT-6 secretion machinery protein EssA |
| SA0275 | 0.606766 | EssB | Type VII secretion system protein EssB |
| SA0276 | 0.516419 | EssC | Type VII secretion system protein EssC |
| SA0272 | 0.56005 | EsaA | Type VII secretion system accessory factor EsaA |
| [SA1862](https://www.genome.jp/entry/sau:SA1862) | -0.84772 | LeuA | 2-isopropylmalate synthase |
| [SA1863](https://www.genome.jp/entry/sau:SA1863) | -0.81808 | LeuB | 3-isopropylmalate dehydrogenase |
| [SA1864](https://www.genome.jp/entry/sau:SA1864) | -0.69053 | LeuC | 3-isopropylmalate/(R)-2-methylmalate dehydratase large subunit |
| [SA1865](https://www.genome.jp/entry/sau:SA1865) | -0.95587 | LeuD | 3-isopropylmalate/(R)-2-methylmalate dehydratase small subunit |
| [SA1866](https://www.genome.jp/entry/sau:SA1866) | -0.94255 | IlvA | threonine dehydratase |
| [SA1859](https://www.genome.jp/entry/sau:SA1859) | -0.74303 | IlvB | E2.2.1.6L; acetolactate synthase I/II/III large subunit |
| [SA1861](https://www.genome.jp/entry/sau:SA1861) | -0.75075 | IlvC | alpha-keto-beta-hydroxylacil reductoisomerase |
| [SA2435](https://www.genome.jp/entry/sau:SA2435) | -0.75769 | ManA | mannose-6-phosphate isomerase |
| [SA2434](https://www.genome.jp/entry/sau:SA2434) | -0.7338 | ManP | mannose PTS system EIIBCA component |
| [SA1962](https://www.genome.jp/entry/sau:SA1962) | -0.62255 | MtlA | mannitol PTS system EIICBA or EIICB component |
| [SA1963](https://www.genome.jp/entry/sau:SA1963) | -0.68691 | MtlD | mannitol-1-phosphate 5-dehydrogenase |
| [SA1960](https://www.genome.jp/entry/sau:SA1960) | -0.79557 | MtlF | PTS system, mannitol specific IIBC component |
| SA0793 | -0.569972307 | DltA | D-alanine--D-alanyl carrier protein ligase |
| SA0794 | -0.547773512 | DltB | Teichoic acid D-alanyltransferase |
| SA0795 | -0.527286241 | DltD | D-alanyl carrier protein ligase |
