## Supplementary Table 3 for "Universal Receptive System as a novel regulator of transcriptomic activity of *Staphylococcus aureus*"

List of differentially expressed genes in *S. aureus* after RNase treatment used for KEGG analysis.

| **Locus** | **Log 2 Fold change** | **Common name** | **Description** |
| --- | --- | --- | --- |
| SA0223 | 2.347171784 | FadA | Acetoacetyl-CoA thiolase |
| SA1037 | 1.61045325 | CatE | VOC domain-containing protein |
| SA0224 | 2.323580088 | FadN | 6-phosphogluconate dehydrogenase |
| SA0225 | 2.057344603 | SA0225 | glutaryl-CoA dehydrogenase |
| SA0534 | 2.5190275 | VraB | Vancomycin resistance-associated protein B |
| SA1418 | 3.543656473 | ComEA | Competence protein EA |
| SA1416 | 2.942128176 | ComEC | Competence protein EC |
| SA0705 | 2.484581377 | ComFA | Competence protein FA |
| SA1374 | 2.563379372 | ComGA | Competence protein GA |
| SA1370 | 4.49715463 | ComGE | Late competence protein ComGE |
| SA1373 | 3.236582814 | ComGB | Competence protein GB |
| SA1372 | 3.589272832 | ComGC | Competence protein G |
| SA1371 | 4.145083662 | ComGD | Competence protein GD |
| SA1486 | 5.914373735 | ComC | Competence protein C |
| SA1369 | 4.136055938 | ComYC | Competence protein YC |
| SA1221 | 2.628792717 | PstS | Phosphate-binding protein PstS |
| SA0068 | 3.314208483 | KdpA | Potassium-transporting ATPase potassium-binding subunit 2 |
| SA1880 | 2.269184773 | KdpB | Potassium-transporting ATPase B chain |
| SA1701 | 2.778256788 | VraS | Sensor histidine kinase VraS |
| SA1700 | 2.376510893 | VraR | Response regulator VraR |
|  |  | VraB | Acetoin dehydrogenase E1 component |
| SA0759_02255 | 2.12024242 | VraD | Acetoin dehydrogenase E2 component |
| SA0536 | 2.389639429 | VraX | VraX protein |
| SA0535 | 3.348576924 | VraC | Acetoin dehydrogenase E3 component |
| SA0759_03489 | 2.569824249 | VraE | VraE protein |
| SA0252 | -2.223962922 | lrgA | Anti-holin-like protein LrgA |
| SA0253 | -2.711698682 | lrgB | Anti-holin-like protein LrgB |
| SA1219 | 2.085466053 | PstA | Phosphate transport system permease protein PstA |
| SA1220 | 2.711698682 | PstC | Phosphate transport system permease protein |
| SA1221 | 2.628792717 | PstS | Phosphate-binding protein PstS |
| SA1218 | 2.188065649 | PstB |  |
|  | 2.188065649 | PhnD | phosphonate transport system substrate-binding protein |
| SA1688 | 2.881661967 | TagH | teichoic acid transport system ATP-binding protein |
| - | 2.785280017 | YydJ | putative peptide transport system permease protein |
| - |  | MntC | manganese transport system substrate-binding protein |
