## Supplementary Table 4 for "Universal Receptive System as a novel regulator of transcriptomic activity of *Staphylococcus aureus*"

List of differentially expressed genes in *S. aureus* after DNase and RNase treatment used for KEGG analysis.

| **Locus** | **Log 2 Fold change** | **Common name** | **Description** |
| --- | --- | --- | --- |
| SA2291 | 0.720554193 | fnbA | Fibronectin-binding protein |
| SA2290 | 0.824457533 | fnbB | Fibronectin-binding protein B |
| SA0521 | 0.677357201 | sdrD | Serine-aspartate repeat-containing protein D |
| SA0107 | 0.549650502 | spa | Immunoglobulin G-binding protein A |
| SA0272 | 0.855038659 | esaA | Type VII secretion system accessory factor EsaA |
| SA0273 | 0.889200772 | essA | ESAT-6 secretion machinery protein EssA |
| SA0275 | 0.906011452 | essB | Type VII secretion system protein EssB |
| SA0276 | 1.005592893 | essC | Type VII secretion system protein EssC |
| SA0277 | 0.834040802 | ecxC | ESAT-6 secretion system extracellular protein C |
| SA1862 | -1.05624244 | leuA | 2-isopropylmalate synthase |
| SA1863 | -0.877887876 | leuB | 3-isopropylmalate dehydrogenase |
| SA1864 | -1.072094729 | leuC | 3-isopropylmalate dehydratase large subunit |
| SA1866 | -1.043536363 | ilvA | L-threonine dehydratase biosynthetic IlvA |
| SA1861 | -0.779267339 | IlvC | Ketol-acid reductoisomerase (NADP(+)) |
